# Group A *Streptococcus* antibiotic tolerance in necrotizing fasciitis

**DOI:** 10.1101/2022.08.19.504414

**Authors:** Nadia Keller, Mathilde Boumasmoud, Federica Andreoni, Andrea Tarnutzer, Manuela von Matt, Thomas C. Scheier, Alejandro Gómez-Mejia, Markus Huemer, Ewerton Marques-Maggio, Reto A. Schuepbach, Srikanth Mairpady-Shambat, Silvio D. Brugger, Annelies S. Zinkernagel

**Author notes:** These authors contributed equally (Nadia Keller, Mathilde Boumasmoud, Federica Andreoni). Corresponding author **Corresponding author contact information** Prof. Annelies S. Zinkernagel, MD PhD.

## Abstract

**Objectives:** Group A *Streptococcus* (GAS) necrotizing fasciitis (NF) is a difficult-to-treat bacterial infection associated with high morbidity and mortality despite extensive surgery and targeted antibiotic treatment. Bacteria surviving prolonged antibiotic exposure without displaying genetic resistance, so-called persisters, are associated with difficult-to-treat infections, such as GAS-NF. In the present study, we investigated the presence of persistent GAS in tissue freshly debrided from three NF patients and examined more in depth the persisters phenomenon in GAS-NF clinical isolates.

**Methods:** Time-lapse imaging of freshly isolated GAS-NF clinical isolates, the image analysis software ColTapp and antibiotic challenge-based persisters assays were used to assess the presence of persisters.

**Results:** We show for the first time that GAS recovered directly from freshly debrided NF patient’s tissue are characterized by increased colony appearance time heterogeneity, indicating the presence of persisters. Acidic pH or nutrient stress exposure, mimicking the NF-like environment in vitro, similarly leads to phenotypic heterogeneity resulting in enhanced antibiotic survival and confirming the presence of GAS persisters.

**Conclusions:** GAS persisters are present in the tissue freshly debrided from GAS-NF patients and might be one explanation for antibiotic treatment failure and surgery requirement in GAS-NF. Tailored treatment options, including the use of persisters-targeting drugs, need to be developed to increase GAS-NF therapy success.

## Background

Necrotizing fasciitis (NF) caused by the human pathogen Group A *Streptococcus* (GAS, *aka Streptococcus pyogenes*) remains a devastating disease, associated with high morbidity and mortality despite the use of effective antibiotics and surgery [1]. Bacteria employ several strategies to escape killing, including antibiotic resistance development, intracellular localization and biofilm formation, the last two protecting them from antibiotic exposure and the immune system [2]. Antibiotic tolerance describes the ability of a fully antibiotic-susceptible bacterial population to survive antibiotic exposure for a prolonged time [3], while antibiotic persistence describes a bacterial subpopulation displaying such ability. Individual surviving bacteria are called persisters and are characterized by an altered metabolism allowing them to withstand adverse conditions. Once the stress is removed, persisters can thrive again and cause relapsing infections. Persistence has been extensively studied in Gram-negative as well as, more recently, Gram-positive bacteria [4, 5]. Persisters are difficult to identify in patient samples. Colony appearance time, which reflects time until bacterial cell growth resumption upon plating, has been used as a proxy to quantify persisters in clinical samples [6, 7].

This fuels the question whether an antibiotic tolerant bacterial sub-population might lead to empiric treatment failure in GAS NF. The presence and nature of persisters in invasive GAS infections has not been investigated so far. We therefore monitored colony growth of GAS directly isolated from NF patients. Moreover, we quantified survival of the NF clinical isolates in vitro upon pre-exposure to acidic pH stress followed by antibiotic-challenge. Here we describe for the first time the persisters phenomenon in GAS derived directly from NF patients, adding another possible explanation for antibiotic treatment failure in NF.

## Methods

### Patient information and samples processing

Three patients presenting with GAS-NF at the University Hospital Zurich and for whom tissue samples were obtained after surgery, were enrolled in this study between 2017 and 2019. Patient parameters, including clinical isolates information and treatment regimens are depicted in Tab.1. Patient samples were freshly processed immediately after surgery for isolation of bacteria and monitoring of colony appearance time. Tissue samples were homogenized in PBS in a tissue lyser (Qiagen). After centrifugation the supernatant was collected and the pellet washed twice with PBS and resuspended in deionized sterile water. Serial dilutions were spread on blood agar plates (Columbia+5% sheep blood, BioMérieux) for time-lapse imaging. Patient tissues were processed as previously described for histological evaluation [8].

**Table 1.**
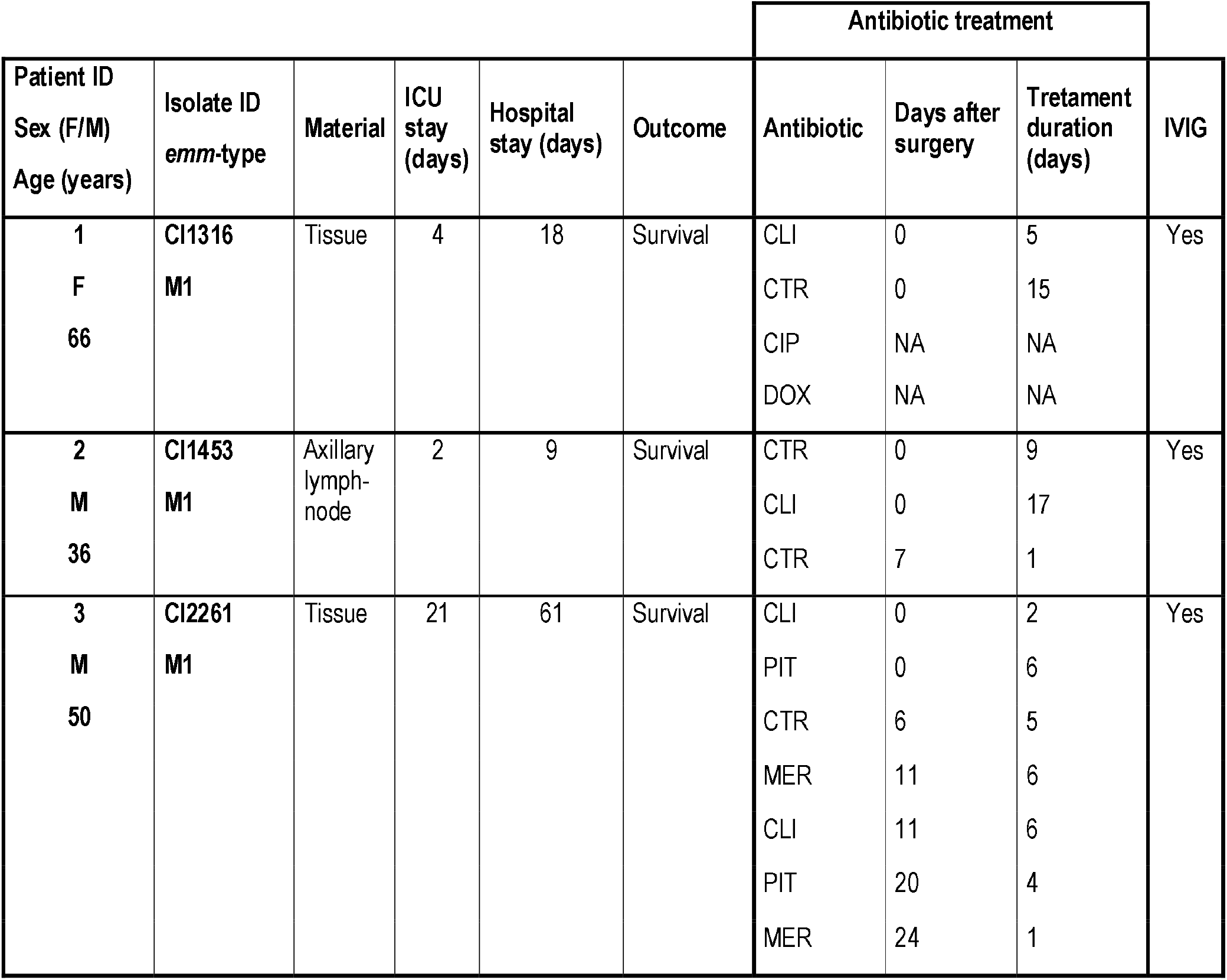
Patient information. Patient and strain information is depicted in panel A. If antibiotic therapy was started before surgery, the symbol “0” is depicted in the column “days after surgery”. Abbreviations: AMC=amoxicillin/clavulanic acid, CIP=ciprofloxacin, CLI=clindamycin, CTR=ceftriaxone, CUR=cefuroxime, DOX=doxycycline, MER=meropenem, PIT=piperacillin/tazobactam, NA=not available, IVIG=intravenous immunoglobulins.

| Patient ID<br>Sex (F/M)<br>Age (years) | Isolate ID<br>*emm*-type | Material | ICU<br>stay<br>(days) | Hospital<br>stay (days) | Outcome | Antibiotic treatment |  |  | IVIg |
| --- | --- | --- | --- | --- | --- | --- | --- | --- | --- |
|  |  |  |  |  |  | Antibiotic | Days after<br>surgery | Treatment<br>duration<br>(days) |  |
| 1<br>F<br>66 | CI1316<br>M1 | Tissue | 4 | 18 | Survival | CLI | 0 | 5 | Yes |
|  |  |  |  |  |  | CTR | 0 | 15 |  |
|  |  |  |  |  |  | CIP | NA | NA |  |
|  |  |  |  |  |  | DOX | NA | NA |  |
| 2<br>M<br>36 | CI1453<br>M1 | Axillary<br>lymph-<br>node | 2 | 9 | Survival | CTR | 0 | 9 | Yes |
|  |  |  |  |  |  | CLI | 0 | 17 |  |
|  |  |  |  |  |  | CTR | 7 | 1 |  |
| 3<br>M<br>50 | CI2261<br>M1 | Tissue | 21 | 61 | Survival | CLI | 0 | 2 | Yes |
|  |  |  |  |  |  | PIT | 0 | 6 |  |
|  |  |  |  |  |  | CTR | 6 | 5 |  |
|  |  |  |  |  |  | MER | 11 | 6 |  |
|  |  |  |  |  |  | CLI | 11 | 6 |  |
|  |  |  |  |  |  | PIT | 20 | 4 |  |
|  |  |  |  |  |  | MER | 24 | 1 |  |

### Colony imaging and appearance time definition

Colony growth was monitored with an automated imaging system, acquiring pictures of the blood agar plates at 10 minutes intervals [9]. Downstream analysis was performed with ColTapp [9]. Colony appearance-time, defined as the time to reach a radius of 200 µm, was directly derived from the colony radial growth curves.

### Bacterial growth conditions

GAS strains M1T1 5448 (GAS WT) [10] and its animal passaged version (GAS AP) carrying a mutation in the *covS* gene, leading to upregulation of virulence factors expression [11] and the three clinical isolates were cultivated in Todd Hewitt Broth (THB; BD) supplemented with 2% yeast extract (THY; Oxoid) and grown at 37°C in a static incubator. CI1316, CI1453 and CI2261 strains were isolated at the University Hospital Zurich from NF patient material (Tab.1). For acidic pH stress, bacteria were grown overnight (O/N) on blood agar plates, resuspended in pH 6 medium [60% THY+30% H2O+10% pH 4.6 buffer (46.8% Na2HPO4 0.2M+53.2% citric acid 0.1M)] to an optical density at 600 nm (OD600) of either 0.2 for GAS WT, CI1453 and CI2261 or 2.0 for GAS AP and CI1326 and incubated at 37ºC for 24h. For neutral pH exposure, bacteria were grown O/N on blood agar plates, resuspended in pH 7.4 medium [60% THY+40% pH 7.5 buffer (90.85% Na2HPO4 0.2M+9.15% citric acid 0.1M)] to an OD600 of 0.2 and incubated at 37ºC for 24h. Serial dilutions of all cultures were subsequently plated for colony enumeration (THY agar) and monitoring of colony appearance time (blood agar). Survival was calculated based on the initial inoculum. Exponential growth phase bacteria were obtained by diluting O/N cultures 1:100 in fresh THY and allowing regrowth to an OD600 of 0.4. Stationary growth phase bacteria were sampled directly from a blood plate after O/N growth.

### Antibiotic susceptibility evaluation

The minimal inhibitory concentration (MIC) of ceftriaxone was determined with the broth microdilution assay in THY medium. The MIC was 0.0625 ug/ml for all strains.

### Persister assay

To determine the fraction of persisters in each condition (exponential or stationary growth phase and acidic pH stress exposure), 5*10^7^colony forming units (CFUs)/ml were challenged with 40x MIC ceftriaxone in THY medium at 37ºC for 24h. Bacteria were washed twice with PBS and serial dilutions plated to assess survival.

### Genotyping

Genotypic characterization of the clinical isolates included typing of the *emm* gene, encoding the surface M-protein, and of the *covS* gene, involved in virulence factors expression regulation. Typing of the *emm* and *covS* genes was performed in silico, based on the whole-genome of the five GAS strains sequenced, assembled and annotated as previously described [12]. In brief, GAS strains were whole-genome sequenced on a MiSeq instrument. De novo-assemblies were built with SPAdes v.3.10 [13] with the --careful command and default k-mer size. The assemblies were annotated with Prokka and the CovS protein sequence was extracted from the .faa file. Variant calling across isolates was performed with Snippy v.4.4.3 [14], by aligning the *trimmed* reads to the de *novo* assembly of one of the isolates as well as to the .gbk file of MGAS5005 (Genbank accession: NC_007297) to assess the effects of variants. The *emm* gene was typed in silico using Blastn 2.9.0 and the multi-fasta file containing the trimmed EMM variant sequences available in the CDC database in August 2021 [15]. All Illumina paired-end reads are available through the European Nucleotide Archive project PRJEB52606.

### Statistics

To determine the effect of clinical isolate and condition on median colony appearance-time, a two-way ANOVA was performed using the anova function in R 4.1.3. Subsequently, specific pairwise comparisons were computed by estimate marginal post-hoc tests (multivariate t-distribution based p-value correction, emmeans package). To analyze statistical significance of survival upon antibiotic challenge, the Mann-Whitney test was used (GraphPad).

### Informed consent and ethical approval

Samples and data were collected at the University Hospital Zurich in accordance with the declaration of Helsinki and with the approval of the Canton’s Ethical Committee (Kantonale Ethikkommission Zurich, Switzerland, BASEC 2016-00145, BASEC 2017-02225 and BASEC 2019-01735).

## Results

### Patients and clinical isolates

Three patients with GAS-NF were enrolled in this study and the three corresponding GAS isolates were retrieved for analysis. Two out of three patients were male, their median age was 50 years and the median hospital stay was 18 days (Tab.1). All patients received the protein synthesis inhibitor clindamycin and a β-lactam antibiotic immediately after hospitalization, as well as intravenous immunoglobulin (Tab.1). Histological analysis showed inflammation and infiltration of immune cells, mainly neutrophils, into the tissue (Fig.1A), fibrin clots (Fig.1A top panels) and bacteria in clusters (Fig.1B top panels) as well as engulfed in immune cells (Fig.1B bottom panels). All isolates were of M-type 1 (M1). CI1316 harbored a mutation in the *covS* gene, resulting in a truncated *CovS* protein (Supplementary figure 1).

**Figure 1.**
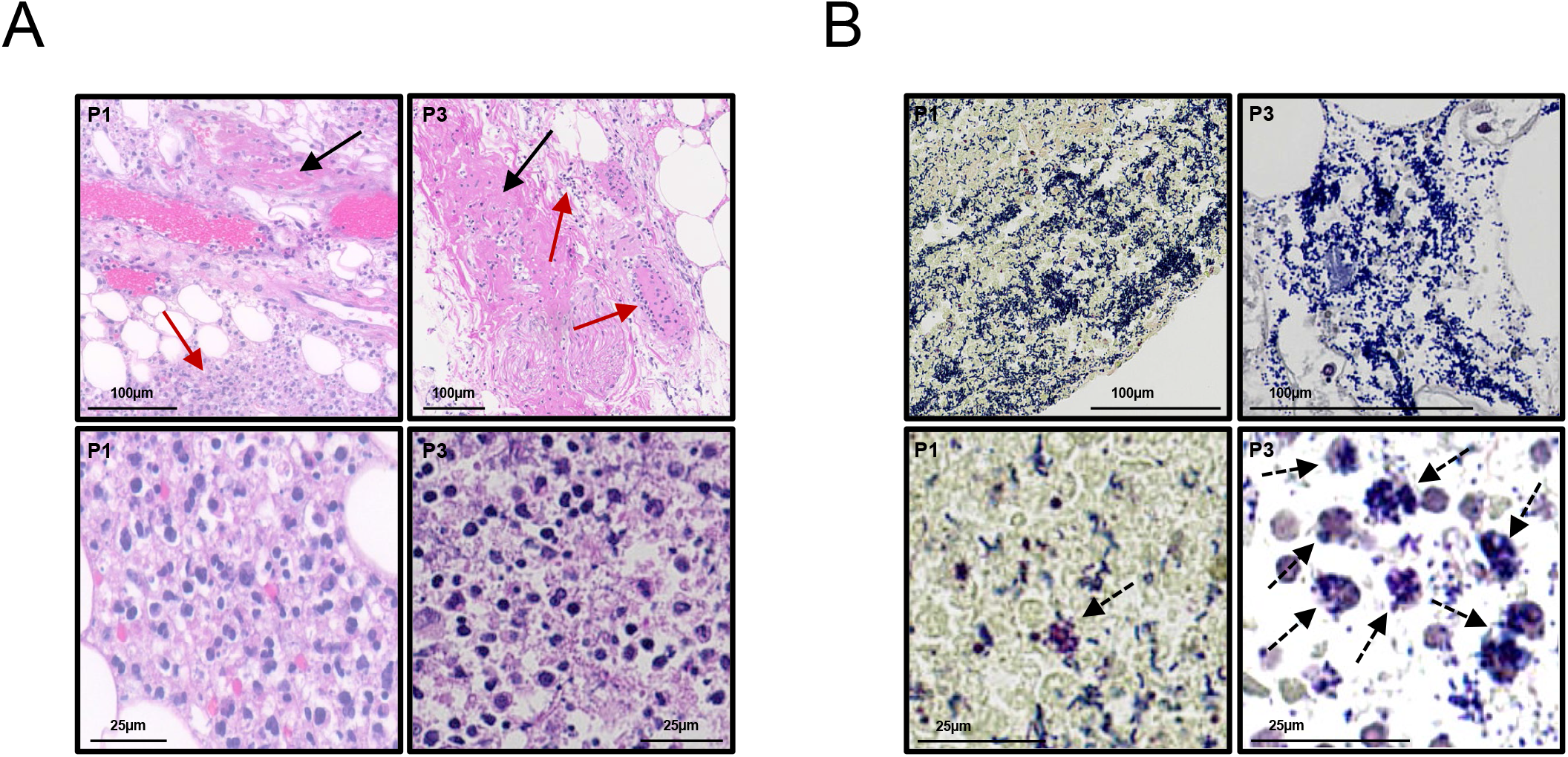
Histology. **A)** Hematoxylin-Eosin stain. Upper panels: the black arrows indicate fibrin clots; the red arrows indicate inflammatory infiltrates. Lower panels: inflammatory infiltrates mainly constituted by neutrophils. B) Brown-Brenn stain. Upper panels: dense biofilm-like bacterial aggregates present in the tissue. Lower panels: the dashed arrows indicate intracellularly located GAS. P1=patient 1; P3=patient 3

### Delayed colony appearance time in GAS isolated directly from patients upon surgery

GAS strains isolated from patient tissue immediately after surgery were plated on blood agar and colony appearance time was monitored. To mimic the clinical situation in vitro, bacteria were pre-exposed to acidic pH or nutrient stress for 24h and subsequently plated, as were exponential growth phase bacteria. Exponential growth phase bacteria (exp) formed detectable colonies after 10h of growth (Fig.2A). Bacteria recovered directly from patient samples (patient) as well as bacteria pre-exposed to acidic pH (THY_pH6) or neutral pH (THY_pH7, nutrient starvation) for 24h, mimicking the in vivo environment, displayed delayed colony appearance times as compared to the exponential growth phase condition (Fig.2A). Bacteria exposed to nutrient starvation and acidic pH survived less than their counterparts exposed to nutrient starvation in the absence of pH stress (Supplementary figure 2). A mutated covS gene, commonly occurring in GAS strains isolated from invasive diseases and leading to upregulation of several virulence factors, did not influence the colony appearance time (Fig.2A).

**Figure 2.**
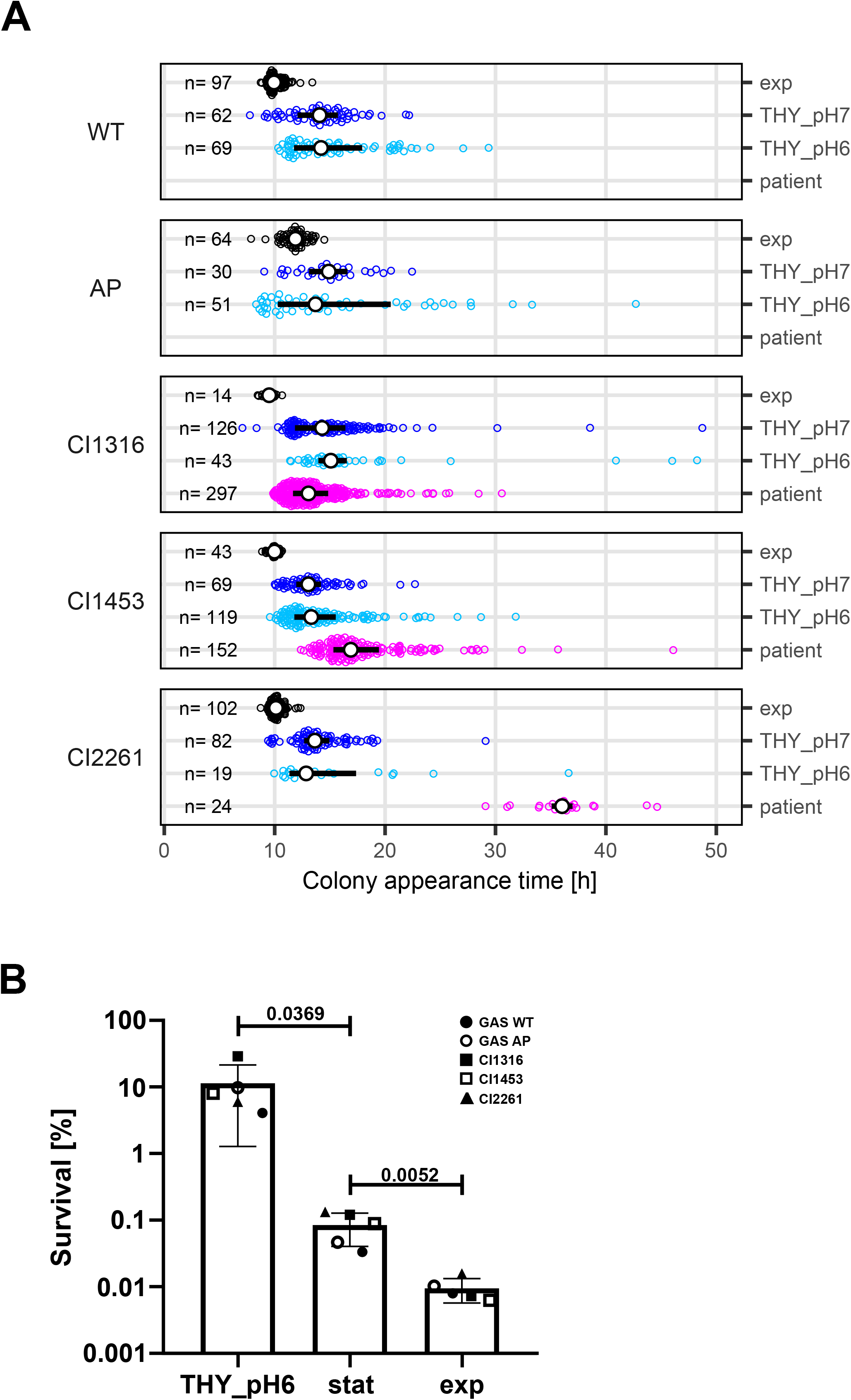
Clinical isolates lag-time and antibiotic tolerance. **A)** Colony appearance time distributions of GAS WT, GAS AP and the three clinical isolates derived from either exponential growth phase (exp) or stationary cultures exposed to acidic or neutral pH (THY_pH6, THY_pH7) are shown. Additionally, for the three clinical isolates, the distribution obtained directly upon sampling from the patient is shown (patient). The number of colonies on each plate is shown on the left (n). The interquartile range and median of each distribution are depicted with a black line and a white dot respectively. A two-way ANOVA revealed that while “isolate” did not have a significant effect on median appearance-time (p-value= 0.59), “condition” had a borderline significant effect (p-value=0.054). Subsequently, estimated marginal means post-hoc tests were performed to compare the median appearance-time averaged across isolates of exponential cultures against other conditions (exp vs.THY_pH7, p-value=0.53; exp vs. THY_pH6, p-value=0.56; exp vs. patient p-value=0.024; p-value corrections based on multivariate t-distribution). B) Antibiotic survival of the various strains after exposure to acidic pH stress for 24 hours (THY_pH6) and sampled either from stationary growth phase cultures (stat) or from exponential growth phase cultures (exp) in THY medium. Each data point represents the average of at least three biological replicates. Statistical analysis was carried out using the Mann-Whitney test. P-values are indicated on the graph.

### Acidic pH pre-exposure and nutrient limitation enhance antibiotic tolerance

To verify whether the populations delayed in growth resumption obtained upon in vitro stress were indeed characterized by an increased antibiotic tolerance, we estimated the baseline survival rate of exponentially growing bacteria upon 24h challenge with 40X MIC of ceftriaxone. Then we exposed bacteria having reached stationary phase to the same challenge. Bacteria in stationary growth phase survived the antibiotic challenge significantly better as compared to their exponential growth phase counterparts (0.1% survival versus 0.01% survival, respectively) (Fig.2B). Acidic pH pre-exposure triggered a dramatically increased survival (∼10%), indicating the presence of a higher number of persisters in the overall bacterial population (Fig.2B).

## Discussion

We showed that GAS cultured immediately after surgical debridement of NF forms persisters, reflected by delayed colony-appearance-times. By mimicking the clinical situation using acidic pH or nutrient stress exposure, we found that GAS persisters are also observed in vitro. The presence of persisters in NF might in part explain antibiotic treatment failure even though GAS remains fully susceptible to penicillin.

The high morbidity and mortality associated with GAS-NF antibiotic treatment failure is likely multifactorial. Poor penetration of antibiotics into the infected necrotic tissue and biofilm-like structures [8, 16] as well as GAS intracellular localization [17] are probable causes. To further elucidate GAS survival in the host, we examined the ability of GAS to form persisters in patients with GAS-NF and mimicked the clinical situation in vitro. We postulated that conditions encountered in the host, such as the acidic pH found in abscesses, necrotic tissue or intracellularly in lysozymes and the presence of antibiotics used for treatment induce or select for a GAS sub-population resilient to antibiotic treatment [2, 18]. The three patients included in this study were treated with high antibiotic doses before surgery, according to current guidelines. Thus, the bacteria were pre-exposed to antibiotics at the time of surgical debridement but remained viable upon plating. Histology showed the presence of dense bacterial aggregates forming biofilm-like structures [8, 16] and intracellular bacteria were detected in all samples. As expected, bacteria freshly recovered from patient tissue showed different degrees of delayed colony appearance time, indicating the presence of dormant cells in the overall bacterial population. Recreating acidic pH stress as well as nutrient starvation in vitro also led to increased heterogeneity in colony appearance time, confirming that pH stress as well as nutrient starvation can trigger the formation of persisters in GAS. This was reflected in increased antibiotic survival rates when acidic pH stress or nutrient starvation were applied before antibiotic challenge as compared to exponential phase growth cultures, as previously observed for S. *aureus* [6].

In order to compare the clinical isolates with well characterized laboratory strains, the laboratory GAS 5448 wild-type strain previously isolated from an invasive infection [8] and its isogenic *covS* mutant GAS AP [11] were used as a benchmark for GAS behavior. In addition, the role of a dysregulation of the two component regulatory *CovRS* system in response to stress and antibiotic challenge was assessed. The presence of a mutation in the covS gene, as found in GAS AP and the clinical isolate CI1316, causes increased expression of several GAS virulence factors [11]. However, a mutated *covS* did not influence colony appearance time nor antibiotic tolerance in this small sample cohort, suggesting that elevated virulence factors activity does not impact persistence in these settings.

In conclusion, we demonstrated that GAS persisters can be isolated directly from patients suffering from NF. In vitro we showed that persister formation is triggered by exposure of bacteria to stressors mimicking the environmental conditions found in the host such as nutrient starvation and acidic pH. Shifting the treatment focus to a regimen that also targets persister cells [5] could help curbing the number of surviving bacteria contributing to reduce GAS-NF treatment failure.

## Footnote page

### Meetings

This work was presented at the EMBO workshop “New approaches to combat antibiotic resistant bacteria” on June 12-16, 2022 in Ascona, Switzerland.

### Authors contribution

NK: Conceptualization, experimental design, acquisition, analysis and interpretation of data.

MB: Experimental design, acquisition, analysis and interpretation of data, writing of the manuscript.

FA: Experimental design, acquisition, analysis and interpretation of data, writing of the manuscript.

AT: Acquisition and analysis of data, critical reading of the manuscript.

MVM: Acquisition and analysis of data, critical reading of the manuscript.

TCS: Analysis of clinical data.

AGM: Acquisition and analysis of data, critical reading of the manuscript.

MH: Acquisition and analysis of data, critical reading of the manuscript.

EMM: Histological evaluation.

RAS: Clinical coordination, funding.

SMS: Experimental design, analysis of data, critical reading of the manuscript.

SDB: Critical reading of the manuscript.

AZ: Conceptualization, writing of the manuscript, funding.

## Figure legends

**Figure S1 - CovS protein sequences alignment**.

Alignment of the CovS protein sequences. Diversion from the GAS WT sequence are marked in light yellow. GAS AP and CI1316 are characterized by truncation of the CovS C-terminal region.

**Figure S2 - Survival of GAS after 24h exposure to pH6 or pH 7.4 medium**.

The GAS strains were grown for 24h in pH6 (THY_pH6) or pH7.4 (THY_pH7) medium at an inoculum of ∼5*10^7^CFUs/ml. Survival was calculated as a percentage of the inoculum. Statistical analysis was carried out using the Mann-Whitney test. P-values are indicated on the graph.

