## Supplemental figure 1 for "Group A *Streptococcus* antibiotic tolerance in necrotizing fasciitis"

### Supplementary figure 1

```
        1 [         .         .         .         .         :         .         .         . 80
WT        MENQKQKQKKYKNSLPKRLSNIFFVLFFCIFSAFTLIAYSSTNYFLLKKEKQSVFQAVNIVRVRLSEVDSNFTLENLAEV
AP        MENQKQKQKKYKNSLPKRLSNIFFVLFFCIFSAFTLIAYSSTNYFLLKKEKQSVFQAVNIVRVRLSEVDSNFTLENLAEV
CI1316    MENQKQKQKKYKNSLPKRLSNIFFVLFFCIFSAFTLIAYSSTNYFLLKKEKQSVFQAVNIVRVRLSEVDSNFTLENLAEV
CI1453    MENQKQKQKKYKNSLPKRLSNIFFVLFFCIFSAFTLIAYSSTNYFLLKKEKQSVFQAVNIVRVRLSEVDSNFTLENLAEV
CI2261    MENQKQKQKKYKNSLPKRLSNIFFVLFFCIFSAFTLIAYSSTNYFLLKKEKQSVFQAVNIVRVRLSEVDSNFTLENLAEV

       81           .         1         .         .         .         .         :         . 160
WT        LYKNDKTHLRIDDRKGSRVIRSERDITNTLDANQDIYVYNIDKQMIFTTDNEESSPGLHGPIGRVYHDHIEDQYRGFSMT
AP        LYKNDKTHLRIDDRKGSRVIRSERDITNTLDANQDIYVYNIDKQMIFTTDNEESSPGLHGPIGRVYHDHIEDQYRGFSMT
CI1316    LYKNDKTHLRIDDRKGSRVIRSERDITNTLDANQDIYVYNIDKQMIFTTDNEESSPGLHGPIGRVYHDHIEDQYRGFSMT
CI1453    LYKNDKTHLRIDDRKGSRVIRSERDITNTLDANQDIYVYNIDKQMIFTTDNEESSPGLHGPIGRVYHDHIEDQYRGFSMT
CI2261    LYKNDKTHLRIDDRKGSRVIRSERDITNTLDANQDIYVYNIDKQMIFTTDNEESSPGLHGPIGRVYHDHIEDQYRGFSMT

      161           .         .         .         2         .         .         .         . 240
WT        QKVYSNRTGKFVGYVQVFHDLGNYYVIRARLLFWLLVVELFGTSLAYLIILITTRRFLKPLHNLHEVMRNISENPNNLNL
AP        QKVYSNRTGKFVGYVQVFHDLGNYYVIRARLLFWLLVVELFGTSLAYLIILITTRRFLKPLHNLHEVMRNISENPNNLNL
CI1316    QKVYSNRTGKFVGYVQVFHDLGNYYVIRARLLFWLLVVELFGTSLAYLIILITTRRFLKPLHNLHEVMRNISENPNNLNL
CI1453    QKVYSNRTGKFVGYVQVFHDLGNYYVIRARLLFWLLVVELFGTSLAYLIILITTRRFLKPLHNLHEVMRNISENPNNLNL
CI2261    QKVYSNRTGKFVGYVQVFHDLGNYYVIRARLLFWLLVVELFGTSLAYLIILITTRRFLKPLHNLHEVMRNISENPNNLNL

      241           :         .         .         .         .         3         .         . 320
WT        RSDISSGDEIEELSVIFDNMLDKLETHTKLQSRFISDVSHELRTPVAIIKGHIGLLQRWGKDDSDILEESLTATAHEADR
AP        RSDISSGDEIEELSVIFDNMLDKLETHTKLQSRFISDVSHELRTPVAIIKGHNWSLTTLG--------------------
CI1316    RSDISSGDEIEELSVIFDNMLDKLETHTKLQSRFISDVSHELRTPVAIIKGHIGLLQRWGKDDSDILEESLTATAHEADR
CI1453    RSDISSGDEIEELSVIFDNMLDKLETHTKLQSRFISDVSHELRTPVAIIKGHIGLLQRWGKDDSDILEESLTATAHEADR
CI2261    RSDISSGDEIEELSVIFDNMLDKLETHTKLQSRFISDVSHELRTPVAIIKGHIGLLQRWGKDDSDILEESLTATAHEADR

      321           .         .         :         .         .         .         .         4 400
WT        MAIMINDMLDMVRVQGSFEGHQNDMTVLEDSIETVVGNFRVLREDFIFTWQSENPKTIARIYKNHFEQALMILIDNAVKY
AP        --------------------------------------------------------------------------------
CI1316    MAIMINDMLDMVRVQGSFEGHQNDMTVLEDSIETVVGNFRVLREDFIFTWQSENPKTIARI-------------------
CI1453    MAIMINDMLDMVRVQGSFEGHQNDMTVLEDSIETVVGNFRVLREDFIFTWQSENPKTIARIYKNHFEQALMILIDNAVKY
CI2261    MAIMINDMLDMVRVQGSFEGHQNDMTVLEDSIETVVGNFRVLREDFIFTWQSENPKTIARIYKNHFEQALMILIDNAVKY

      401           .         .         .         .         :         .         .         . 480
WT        SRKEKKIAINLSVTGKQEAIVRVQDKGEGISKEDIEHIFERFYRTDKSRNRTSTQAGLGIGLSILKQIVDGYHLQMKVES
AP        --------------------------------------------------------------------------------
CI1316    --------------------------------------------------------------------------------
CI1453    SRKEKKIAINLSVTGKQEAIVRVQDKGEGISKEDIEHIFERFYRTDKSRNRTSTQAGLGIGLSILKQIVDGYHLQMKVES
CI2261    SRKEKKIAINLSVTGKQEAIVRVQDKGEGISKEDIEHIFERFYRTDKSRNRTSTQAGLGIGLSILKQIVDGYHLQMKVES

      481           .         ] 500
WT        ELNEGSVFILHIPLAQSKES
AP        --------------------
CI1316    --------------------
CI1453    ELNEGSVFILHIPLAQSKES
CI2261    ELNEGSVFILHIPLAQSKES
```
