## Supplementary figures and images for "Group A *Streptococcus* antibiotic tolerance in necrotizing fasciitis"

### Supplemental figure 2

Supplementary Figure 2

**A**

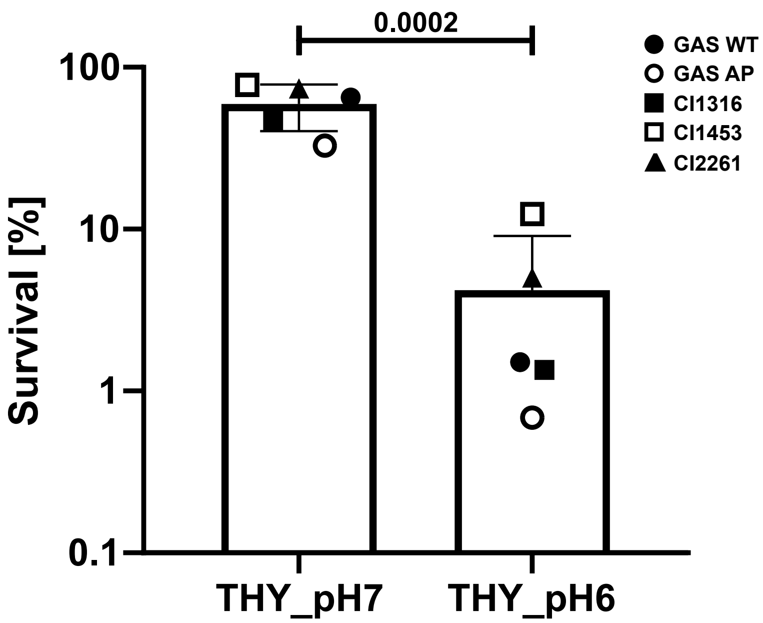
